# Divergent consequences of PSEN1 knockout and PSEN2 knockout in stem cell derived models of the brain

**DOI:** 10.64898/2026.04.09.717238

**Authors:** Charles Arber, Mateo Barro Fernandez, Claudio Llerena Villegas, Lucrezia Bruno, Filip Tomczuk, Patrick A. Lewis, Jennifer M. Pocock, John Hardy, Selina Wray

## Abstract

γ-secretase is a multi-subunit enzyme complex responsible for cleaving hundreds of substrates in diverse cellular contexts. Variation in subunit composition - including the use of alternate catalytic subunits Presenilin 1 (PSEN1) and Presenilin 2 (PSEN2) - results in diverse γ-secretase complexes. Point mutations in *PSEN1* and *PSEN2* cause familial forms of Alzheimer’s disease, while loss-of-function mutations in the γ-secretase subunits *PSEN1, PSENEN* and *NCSTN* cause acne inversa. To advance therapeutic strategies targeting γ-secretase in Alzheimer’s disease, a better understanding of individual γ-secretase complexes is required.

In this study, we used CRISPR-Cas9 genome engineering to generate PSEN2-knockout iPSCs in order to compare the consequence of PSEN2 knockout versus PSEN1 knockout in iPSC-derived brain cells. In contrast to PSEN1-knockout, PSEN2-knockout did not alter APP cleavage or Aβ generation in iPSC-neurons, nor did it disrupt Nicastrin maturation. Similarly, PSEN2-knockout had little impact on TREM2 processing in iPSC-microglia. Instead, our data indicate that loss of PSEN2 primarily impacts the endo-lysosomal system in iPSC-neurons, causing an accumulation of early endosome markers and a reduction in lysosomal markers – phenotypes not observed in PSEN1-knockout neurons.

Taken together, these findings highlight distinct and non-redundant functions of PSEN1 and PSEN2 in human brain cells, reinforcing findings in animal models and subcellular localisation studies. This work advances our understanding of distinct γ-secretase complex functions and provides insights that will support future therapeutic efforts to inhibit, modulate or stabilise γ-secretase.

## Introduction

γ-secretase is a multi-subunit enzyme complex responsible for the regulated intramembrane proteolysis of several hundred transmembrane substrates [1,2]. The complex consists of 4 components: obligate subunits PEN-2 and Nicastrin as well as alternate subunits: either APH1A or APH1B; and either Presenilin 1 (PSEN1) or Presenilin 2 (PSEN2). PSEN1 and PSEN2 represent the catalytic subunits, harbouring aspartate residues responsible for hydrolysis within the catalytic pore [3]. Dominantly inherited mutations in *PSEN1* and *PSEN2* cause familial Alzheimer’s disease (fAD) [4,5], whereas homozygous loss-of-function mutations in *PSEN1, PSENEN* (the gene encoding PEN-2) and *NCSTN* (encoding Nicastrin) cause familial hidradenitis suppurativa and acne inversa [6,7]. γ-secretase cleaves the amyloid precursor protein (APP) as the rate limiting step in the production of amyloid-beta (Aβ), the peptide component of amyloid plaque pathology in Alzheimer’s disease (AD). All known mutations that cause fAD converge on altering the production of Aβ [8].

Alternate subunit usage generates a diverse range of γ-secretase complexes, resulting in functional diversity. For example, the subunit composition can impact the relative catalytic activity of the enzyme [9,10]. γ-secretase complexes have been shown to exhibit different cellular and subcellular enrichment [11]. At the cellular level, PSEN2-associated γ-secretase has been suggested to be enriched in microglia compared to other brain cell types [12], and at the functional level, distinct phosphorylation patterns of PSEN1 and PSEN2 suggest divergent cellular biology [13]. Subcellularly, PSEN1-associated γ-secretase has been shown to be broadly distributed within the cell, whereas PSEN2-enriched γ-secretase is associated with endosomal and lysosomal membranes [14,15].

Previous efforts to inhibit γ-secretase in clinical trials in Alzheimer’s disease failed due to adverse side-effects [16], likely caused by effects on unintended substrates and driven by an incomplete understanding of γ-secretase diversity. A detailed understanding of γ-secretase is essential for developing targeted therapeutic strategies, which aim to modulate γ-secretase [17]. Defining redundant versus unique roles for PSEN1 and PSEN2-associated γ-secretase will be key for subtype-specific targeting and improving the precision of therapeutic strategies. To date, much of our knowledge comes from studies using animal models to investigate Psen1 and Psen2 loss-of-function. *Psen1* knockout mice are embryonic lethal [18,19], a consequence attributed to disrupted Notch signalling in development. In contrast, *Psen2* knockout mice are viable and fertile [20], although they exhibit phenotypes including alveolar wall thickening and mitochondrial dysfunction [21]. When *Psen1* and *Psen2* are both conditionally ablated in postmitotic neurons, their combined loss results in an additive effect, and mice develop progressive neurodegeneration [22]. This underscores their complementary but crucial roles in neuronal homeostasis.

We previously showed that the fAD-associated int4del *PSEN1* mutation and PSEN1 loss-of-function caused distinct cellular phenotypes in isogenic iPSC-neurons [23]. Specifically, PSEN1-knockout caused reduced Nicastrin maturation, an accumulation of APP cleavage fragments, and a 60% reduction in Aβ production: phenotypes distinct from the int4del mutation-harbouring neurons. To extend these findings, and to understand differential consequences of PSEN1 and PSEN2 loss-of-function, we used CRISPR-Cas9 genome editing to generate isogenic PSEN2-knockout iPSC and directly compare the resulting cellular phenotypes to PSEN1-knockout cells.

## Methods

### Genome editing

Isogenic control and PSEN1-knockout iPSCs were generated in a previous study through the correction of the int4del mutation in patient-derived iPSCs [23]. To generate PSEN2-knockout cells, corrected control iPSCs underwent CRISPR-Cas9 genome editing. Predesigned sgRNAs were selected from Integrated DNA Technologies (IDT) Alt-R CRISPR-Cas9 guides. Three guides were compared for cutting efficiency and a single lead guide was selected (Table 1). CRISPR-Cas9 ribonucleoprotein was constructed following the IDT Alt-R protocol, combining tracrRNA, crRNA and S.p. Cas9 nuclease 3NLS. Ribonucleoprotein complexes were transfected into iPSCs using nucleofection (P3 Amaxa, Lonza). Transfected cells were expanded in mTESR (Stem Cell Technologies) plus 10µM Y27632 (Tocris) prior to screening and selected clones were transitioned to Essential 8 media (ThermoFisher). 1 lead clone was characterised in detail, with 2 additional clones used for validation studies. Indels were confirmed by TIDE software [24] and confirmed by ICE software (ICE CRISPR Analysis. 2025. v3.0. EditCo Bio; December 2025). The 5 most likely off-target cut-sites (IDT Off Target predictions) were screened by Sanger sequencing in the lead clone, and no evidence of off-target editing was found. See Table 1 for oligonucleotide sequences used in this study.

**Table 1.**
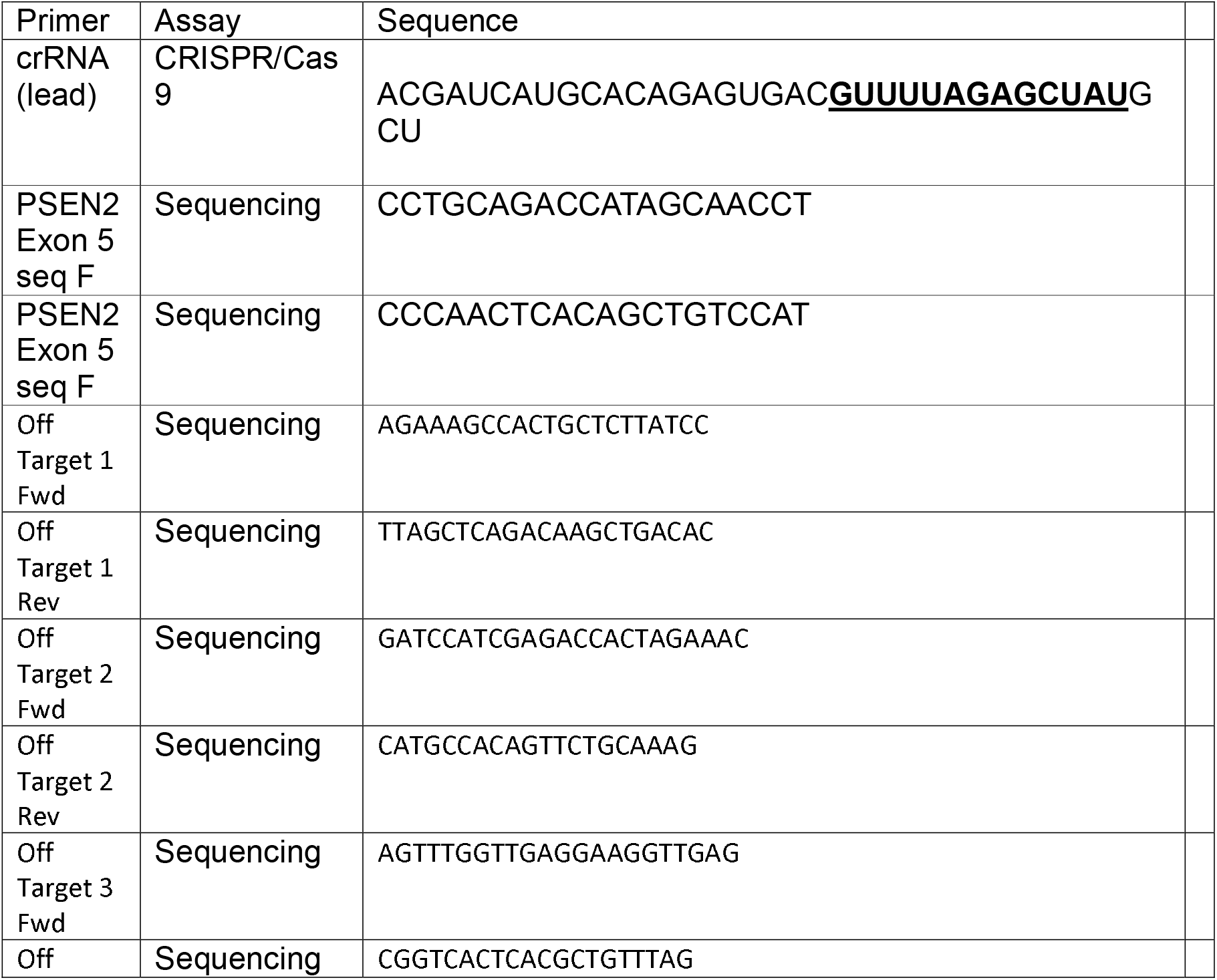
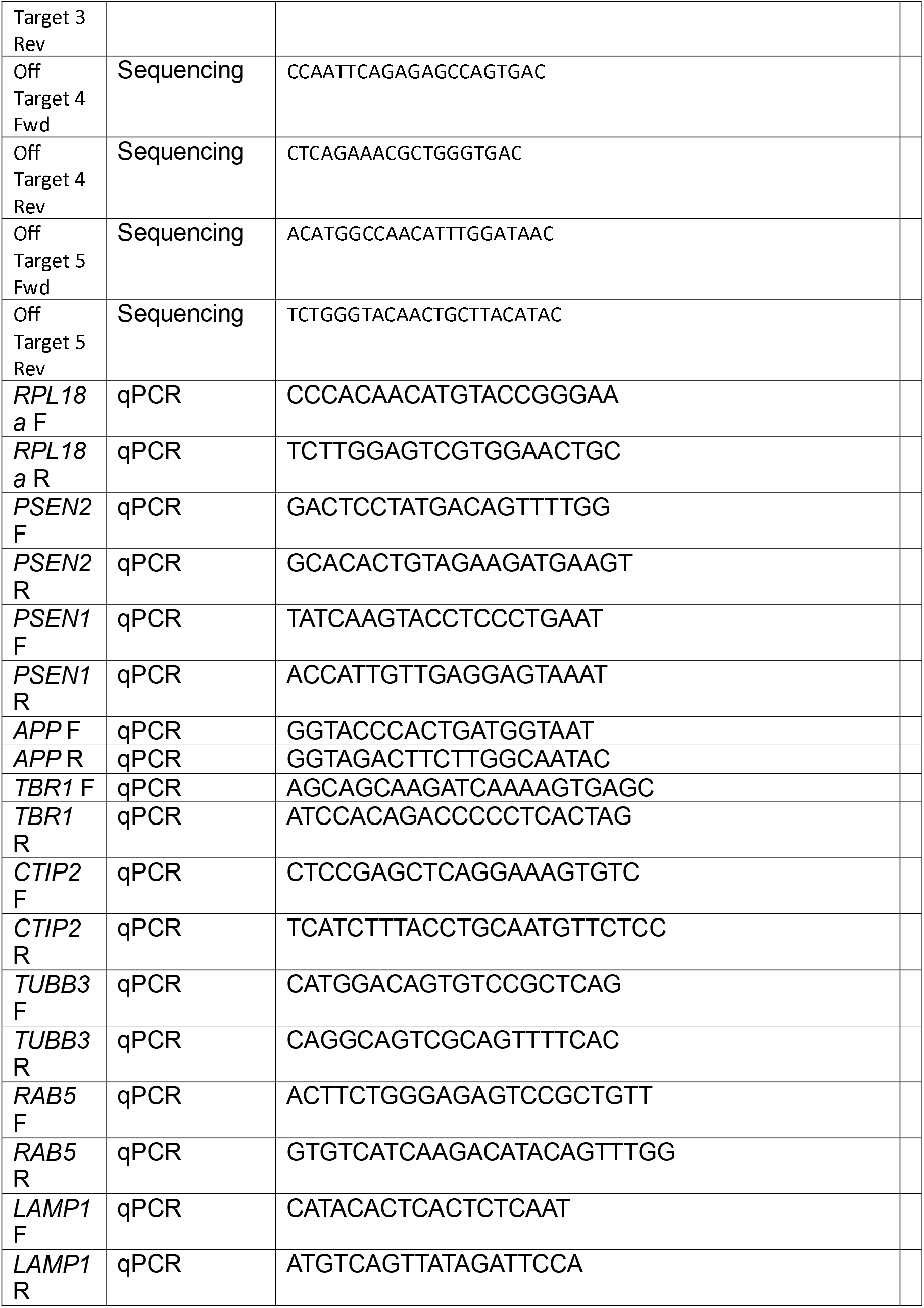
Oligonucleotides used in this study.

### Cell culture

Products are ThermoFisher unless otherwise stated. iPSCs were maintained in Essential 8 media on Geltrex substrate and routinely passaged using 0.5mM EDTA. A stable Karyotype was confirmed using low coverage whole genome sequencing (as previously described [25] and following [26,27]), as well as the Stem Cell Technologies hPSC Genetic Analysis Test (Fig S1).

iPSCs were differentiated to cortical glutamatergic neurons following published protocols [28] previously established [29–31]. Briefly, iPSCs were grown to confluence and media was changed to N2B27 media with 1µM dorsomorphin (Tocris) and 10µM SB431542 (Tocris) for neural induction. N2B27 consists of 50% DMEM-F12, 50% Neurobasal supplemented with 0.5X N2 supplement, 0.5X B27 supplement, 0.5X L-Glutamine, 0.5X non-essential amino acids, 0.5X Pen/Strep, 1:1000 β-mercaptoethanol, and 25U insulin. Cells were passaged onto laminin-coated plates (Sigma), using dispase (DIV10 and DIV18) and accutase (DIV28 and DIV35). Cells were maintained in N2B27 media from day 10 onwards. Day 100 was taken as the final time point for mature cortical glutamatergic neuronal cultures. γ-secretase inhibition was achieved using 10µM DAPT (Tocris) treatment for 48 hours.

iPSCs were differentiated to microglia-like cells (iMGLCs) using published protocols [32,33] previously established [34]. Growth factors were PeproTech unless otherwise stated. Briefly, embryoid bodies were generated consisting of 10,000 iPSCs and cultured in Essential 8 supplemented with 50ng/ml BMP4, 50ng/ml VEGF and 20ng/ml SCF. After day 4, EBs were maintained in X-VIVO 15 media (Lonza) supplemented with 100ng/ml MCSF and 25ng/ml IL3. Maturation consisted of treatment with 100ng/ml IL34, 25ng/ml MCSF, and 5ng/ml TGFβ1 in DMEM-F12 supplemented with 1X N2 supplement, 1X NEAA, 1X L-Glutamine. 1X insulin/selenium/transferrin and 25U insulin. Lipopolysaccharide (Sigma) was used as an inflammatory stimulus at 100ng/ml.

### Western blotting

Samples were lysed in RIPA buffer with protease and phosphatase inhibitors (Roche). The BioRad BCA assay was used to normalise protein content and samples were denatured by boiling in LDS buffer with 20% DTT. Samples were run in precast 4-12% poly-acrylamide gels and transferred to nitrocellulose membranes. Blocking was done in 3% bovine serum albumin, washes were done in 0.1% PBS-Tween-20, and imaging was performed on a Licor Odyssey fluorescent imager. See Table 2 for antibodies used in this study.

**Table 2.**
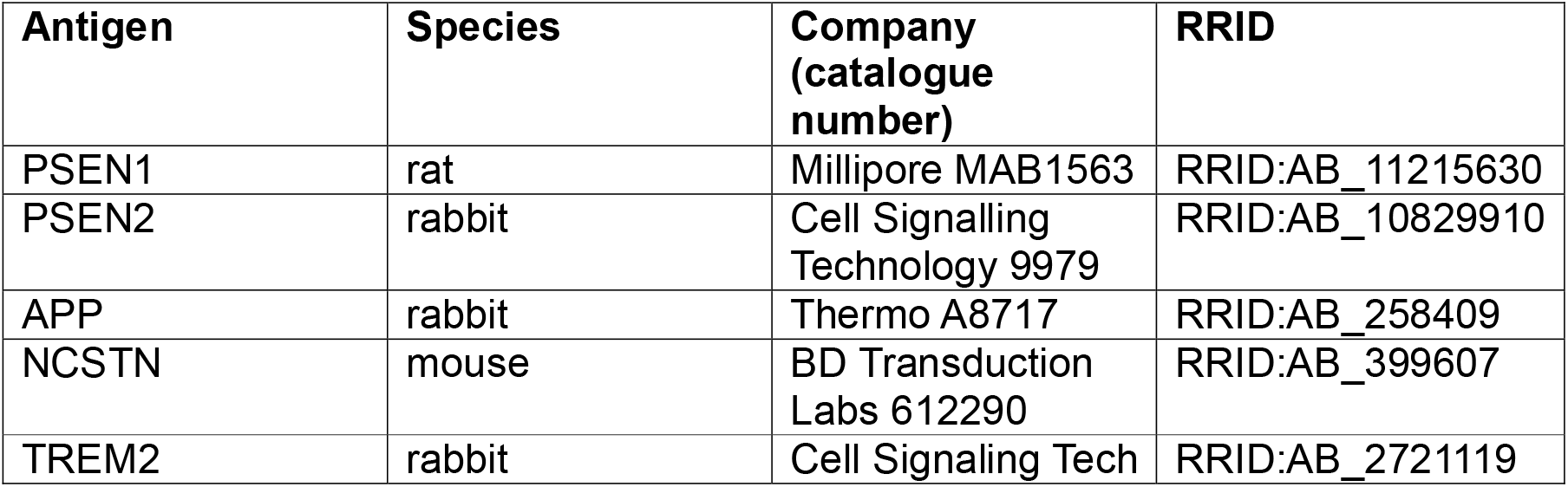
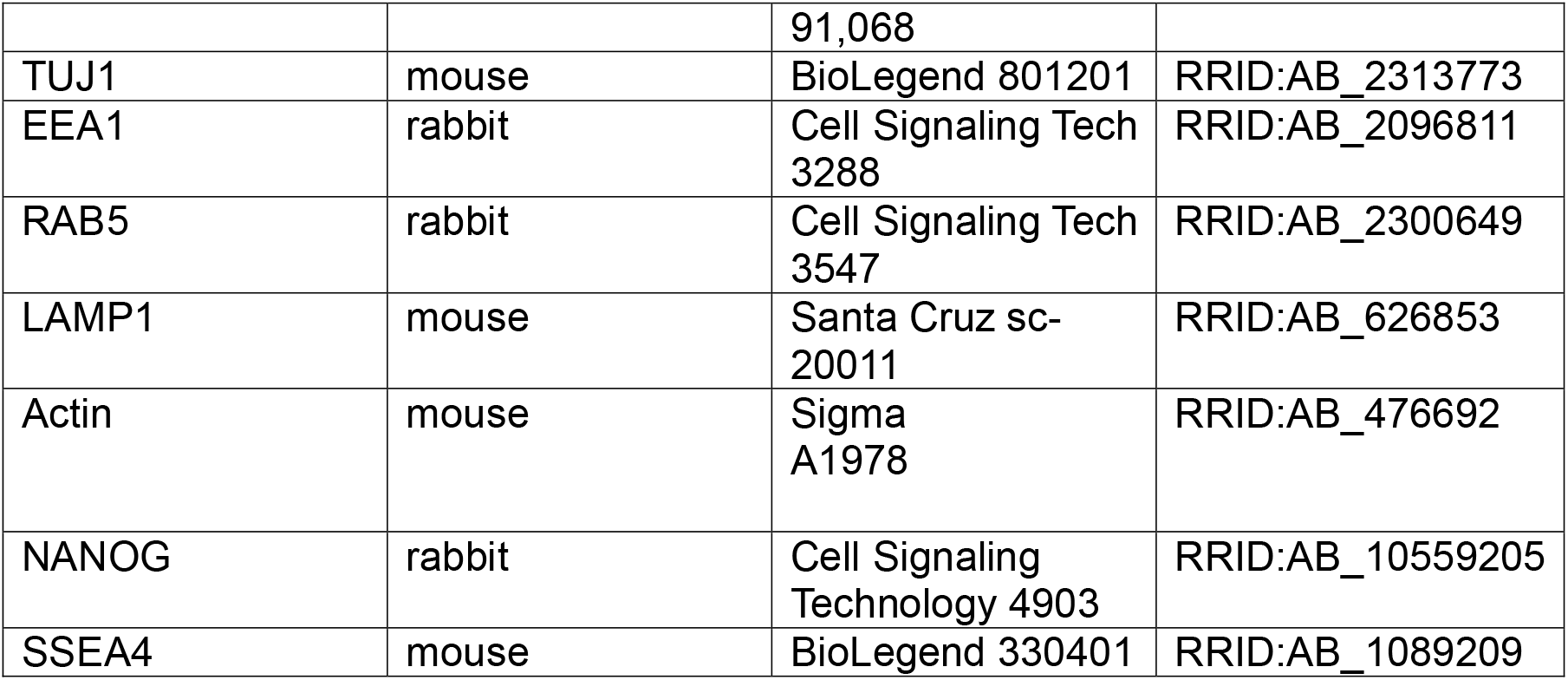
Antibodies used in this study.

### qPCR

Samples were lysed in Trizol and RNA was isolated following manufacturers instructions. 1-2µg of RNA was reverse transcribed using Superscript IV and random hexamers. qPCR was performed using Power Sybr green mastermix on an Agilent Aria MX3000P thermocycler. See Table 1 for primer sequences and all annealing temperatures were 60°C.

### ELISAs

Neuronal conditioned media was conditioned for 48 hours and centrifuged at 2000g for 5 minutes to remove cellular debris. Aβ38, Aβ40, and Aβ42 were quantified using the Meso Scale Discovery V-Plex Aβ Peptide Panel (6E10) on an Meso QuickPlex SQ 120. Aβ was normalised to the protein content of the corresponding cell pellet.

## Statistical analysis

All experiments were done with at least 3 independent differentiations for experimental replication (see Table S1). Data were analysed in Microsoft Excel and GraphPad Prism. Normality testing (D’Agostino) was performed followed by parametric (One Way ANOVA) analysis with multiple comparisons, as described in figure legends.

## Results

### PSEN2-knockout does not affect APP processing and Nicastrin maturation in iPSC-neurons

To investigate the consequence of PSEN2 knockout in human brain cells, we used CRISPR-Cas9 genome engineering to introduce indels into the constitutive exon 5 of *PSEN2*. CRISPR-Cas9 ribonuclear protein complexes were introduced into previously characterised control iPSCs [23] by nucleofection. Clonal iPSC colonies were expanded, and an initial screen was performed via Western blotting for PSEN2 (Fig S1A). One lead clone and two additional clones were taken forward for detailed characterisation. Sanger sequencing revealed biallelic frameshift mutations in *PSEN2* (Fig S1B), confirming loss of function (frameshift mutations were +1/+1, -4/-6 and +1/-8). iPSCs exhibited stable karyotype (Fig S1C), appropriate morphology and expression of pluripotency markers (Fig 1A).

**Figure 1.**
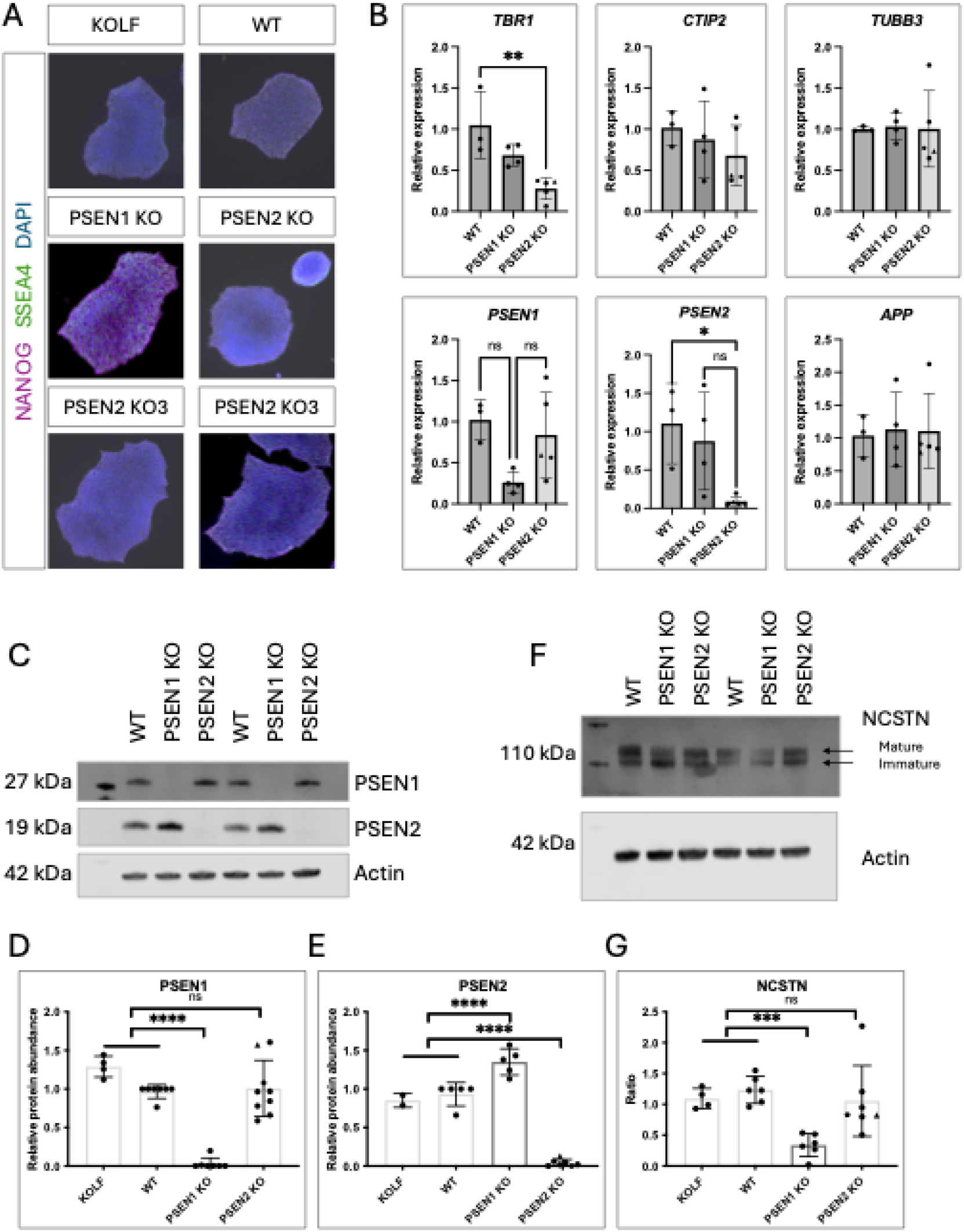
Generation of PSEN2-knockout neurons and the effect on γ-secretase. A) Immunocytochemistry of iPSC lines to show expression of pluripotency markers (NANOG and SSEA4), as well as characteristic iPSC morphology. B) qPCR analysis of isogenic iPSC-derived neurons. Comparable *TUBB3* expression demonstrates similar neuronal content between lines and induction and *TBR1* and *CTIP2* expression represents cortical identity. Data represents at least 3 independent inductions (see Supplementary Table 1); PSEN2 data represents 3 clones (clone 2 represented by hexagon, clone 3 represented by triangle). C) Western blotting confirms absence of PSEN1 and PSEN2 protein in respective knockout cultures. D-E) quantification of PSEN1 and PSEN2 Western blot data. F) Western blotting of Nicastrin maturation (glycosylation). G) Quantification of mature Nicastrin relative to immature Nicastrin. D, E, G) Data represents 4-7 independent inductions (see Supplementary Table 1); PSEN2 data represents 3 clones (clone 2 represented by hexagon, clone 3 represented by triangle). Groups were compared by One Way ANOVA with multiple comparisons. p<0.05 = *, p<0.01 = **, p<0.001 = *** and p<0.0001 = ****.

PSEN2-knockout iPSCs were differentiated into cortical neurons alongside isogenic PSEN1-knockout iPSCs [23] and the unedited parental control line as well as an unrelated, well-characterised control iPSC line, KOLF2.1J [35] (Fig 1B). Loss of PSEN2 expression and protein was demonstrated by qPCR (Fig 1B) and Western blotting (Fig 1C-E). PSEN2 levels were significantly increased in PSEN1-knockout neurons, suggesting a compensatory effect. In contrast, PSEN2 knockout neurons did not show a corresponding increase in PSEN1, suggesting that PSEN1 does not compensate for PSEN2 loss-of-function to the same extent (Fig 1C-E).

One major consequence of PSEN1 loss-of-function is defective glycosylation and maturation of the γ-secretase component Nicastrin [36]. Nicastrin maturation was not impaired in PSEN2 knockout neurons (Fig 1F-G), in contrast to PSEN1-knockout neurons where a reduction in the mature, glycosylated protein is observed.

PSEN1-knockout neurons display an accumulation of APP C-terminal fragments, peptides generated by α- and β-secretase that represent substrates cleaved by γ-secretase to generate Aβ [37]. Analysis of APP cleavage in PSEN2-knockout neurons showed that APP displayed similar processing to control neurons, evidenced via the ratio of full-length APP to its C-terminal fragments (Fig 2A-C). When PSEN1 and PSEN2-knockout neurons were treated with the γ-secretase inhibitor DAPT, further APP C-terminal accumulation was seen, suggesting residual γ-secretase activity in all genotypes (Fig 2D).

**Figure 2.**
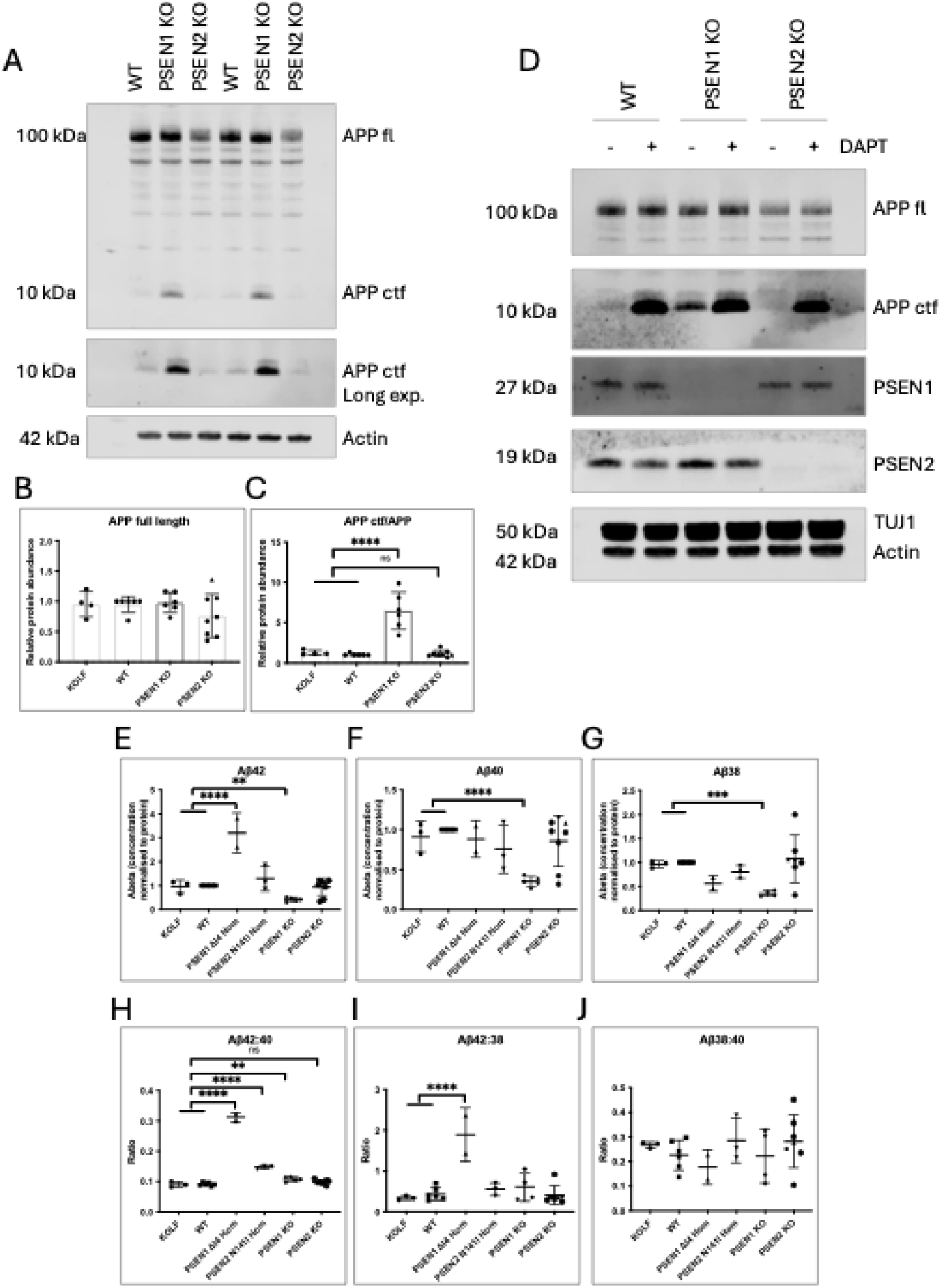
APP processing and Aβ generation is unchanged in PSEN2 knockout neurons. A) Western blotting of APP in iPSC-derived neurons, and B-C) quantification of full-length APP and its C-terminal fragments (CTFs). Data represents 4-6 independent induction (see Supplementary Table 1); PSEN2 data represents 3 clones (clone 2 represented by hexagon, clone 3 represented by triangle). D) Representative Western blot of iPSC-neuronal cultures treated with 10µM DAPT for 48 hours. E-G) Electrochemiluminescent quantification of Aβ42, Aβ40 and Aβ38 normalised to the protein content of each cell pellet. H-J) Ratio data for Aβ quantification. E-J is represented by 3-7 independent replicates (see Supplementary Table 1). Data analysed by one way ANOVA with multiple comparisons. p<0.05 = *, p<0.01 = **, p<0.001 = *** and p<0.0001 = ****.

Aβ generation from PSEN1 and PSEN2-knockout neurons was compared to controls and existing lines homozygous for fAD-associated mutations in *PSEN1* (int4del [23]) and *PSEN2* (N141I [38]). As previously reported, PSEN1-knockout resulted in a reduction of all Aβ species measured (Aβ38, Aβ40 and Aβ42) by around 60-65%, consistent with the accumulation of APP C-terminal fragments (Fig 2 E-G). In contrast, concentration of Aβ peptides did not differ between controls and PSEN2-knockout neuronal cultures (Fig 2E-G). Ratios of Aβ peptides were not significantly different between PSEN2-knockout and control iPSC-neurons (Fig 2H-J), suggesting that loss of PSEN2 does not impair γ-secretase processivity and the generation of Aβ.

Together, these data support divergent effects of PSEN1 and PSEN2 loss-of-function upon APP processing. PSEN2-knockout does not alter APP/Aβ processing or Nicastrin maturation, in contrast to PSEN1 loss-of-function, which results in impaired Nicastrin maturation and reduced APP cleavage to Aβ.

### PSEN2-knockout does not alter TREM2 cleavage in iPSC-derived microglial like cells

PSEN2 has been suggested to act as the major presenilin component of γ-secretase in microglia [12,39], and its expression is higher in iPSC-microglia like cells (iMGLCs) compared with neurons and astrocytes [26]. PSEN1 is also expressed in microglia and has been shown to be crucial for Aβ clearance [40]. TREM2 is a risk factor for AD [41] and is microglial-enriched. It has been shown to be a bona fide γ-secretase substrate [42].

We therefore investigated the impact of PSEN1 and PSEN2-knockout on TREM2 cleavage in iMGLCs under basal conditions and in response to inflammatory stimulation following exposure to lipopolysaccharide. PSEN1-knockout iMGLCs exhibited a significant accumulation of TREM2 C-terminal fragments, the substrate for γ-secretase mediated cleavage after α-secretase shedding of the ectodomain. TREM2 processing was not altered in PSEN2-knockout iMGLCs and relative levels of C-terminal fragments were similar to controls (Fig 3A-B). All genotypes had similar levels of full-length TREM2 at 28kDa with appropriate maturation as shown by higher molecular weight immunoreactivity of post-translationally modified (glycosylated) TREM2 (Fig 3A-B).

**Figure 3.**
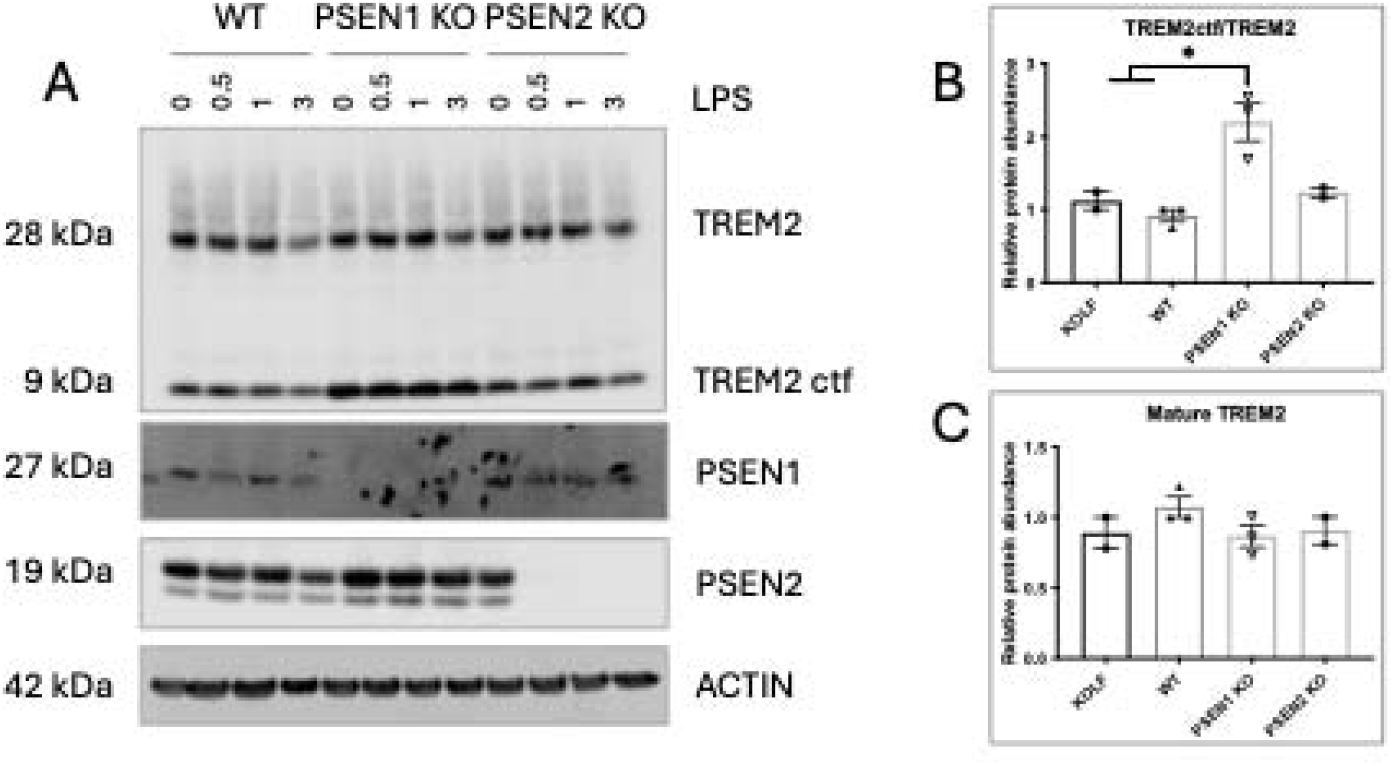
PSEN2-knockout does not alter TREM2 cleavage. A) Western blotting analysis of isogenic iMGLC lysates over a timecourse of LPS treatment (lipopolysaccharide 100ng/ml). B) Quantification of TREM2 levels. Data represents 2-3 independent batches of iMGLCs and up to 3 harvests (see Supplementary Table 1). Groups compared via one way ANOVA with multiple comparisons. p<0.05 = *, p<0.01 = **, p<0.001 = *** and p<0.0001 = ****.

Together these data suggest that PSEN2 loss-of-function does not impair the maturation of TREM2 nor its processing at the plasma membrane in iMGLCs.

### PSEN2-knockout but not PSEN1-knockout affects the endo-lysosomal system in iPSC-neurons

Endo-lysosomal deficits are well described in fAD and PSEN1/2 knockout cells (reviewed in [43]). PSEN1 and PSEN2 have been suggested to be enriched in distinct subcellular localisations [11,14], with PSEN1 being ubiquitous and PSEN2 enriched in endo-lysosomal compartments.

We investigated the consequence of knockout of PSEN1 and PSEN2 on the endo-lysosomal system. Western blotting for early endosome markers EEA1 and RAB5, and lysosomal marker LAMP1 showed no significant alterations in steady state protein levels in PSEN1-knockout neurons. In contrast, PSEN2-knockout neurons displayed a significant accumulation of early endosome markers compared to control iPSC-neurons, and a significant reduction of LAMP1 protein compared to PSEN1-knockout iPSC-neurons (Fig 4 A-E). qPCR analysis supports comparable expression levels of *LAMP1* and *RAB5* between genotypes (Fig 4 F-G), suggesting alterations to endo-lysosomal-associated peptides are post-transcriptional.

**Figure 4.**
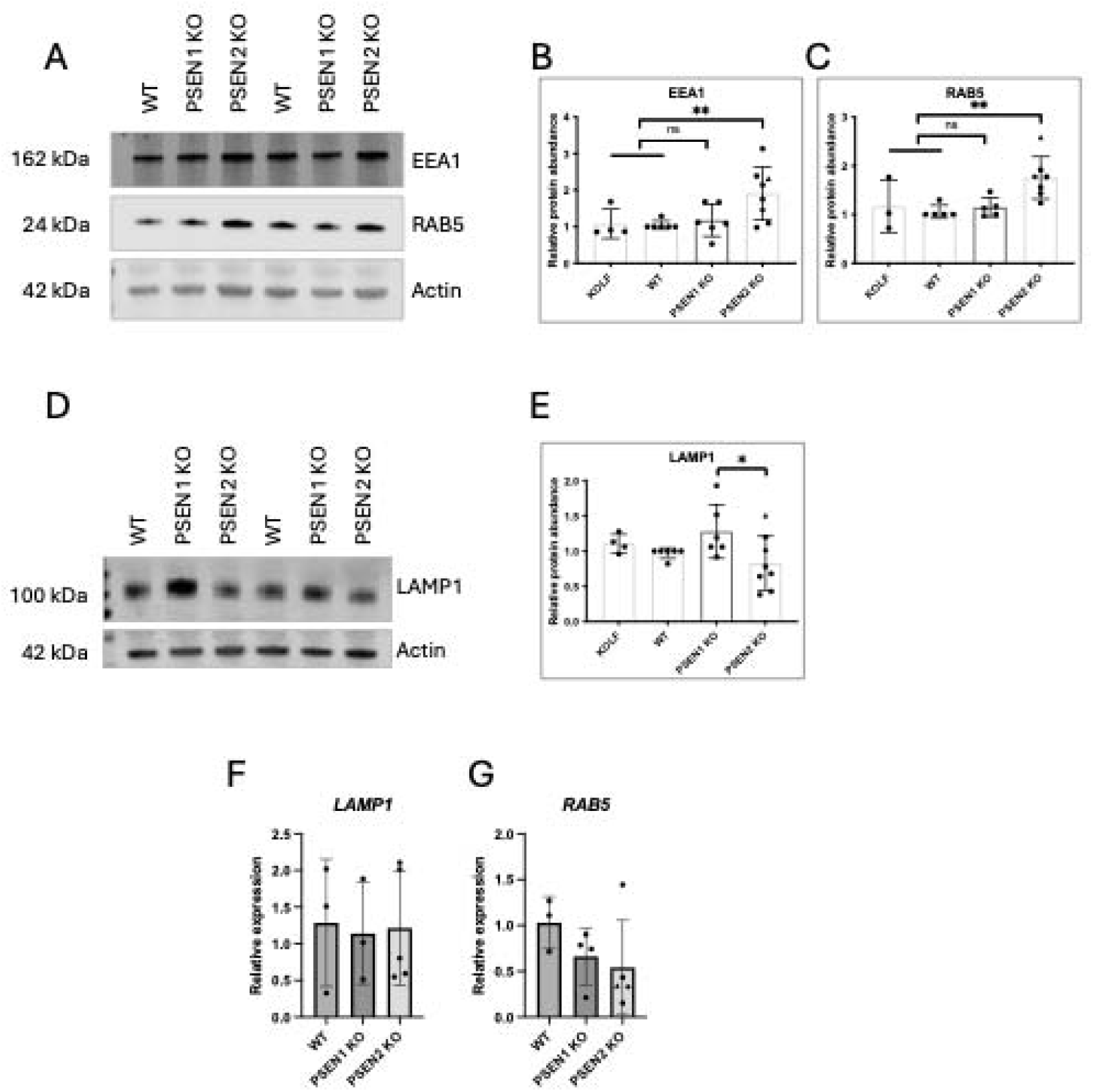
Endo-lysosomal imbalance in PSEN2-knockout iPSC-neurons. A) Western blotting analysis of iPSC-neurons for early endosome markers EEA1 and RAB5. B-C) Quantification of Western blot data in A). D) Western blot analysis of LAMP1 levels in iPSC-neurons, D) quantification. F-G) qPCR analysis for iPSC-neuron expression of lysosome-associated *LAMP1* and endosome-associated *RAB5*. Data represent 3-6 independent repeats (see Supplementary Table 1); PSEN2 data represents 3 clones (clone 2 represented by hexagon, clone 3 represented by triangle). Groups compared via one way ANOVA with multiple comparisons. p<0.05 = *, p<0.01 = **, p<0.001 = *** and p<0.0001 = ****.

Together, these data are suggestive of an impaired endo-lysosomal system associated with PSEN2, but not PSEN1, loss-of-function.

## Discussion

We set out to compare and contrast the consequences of PSEN1 and PSEN2 knockout on substrate processing in neurons and microglia. PSEN2 was effectively knocked out in iPSCs using CRISPR-Cas9 genome editing. PSEN1 and PSEN2-knockout produced distinct effects in iPSC-neurons and iMGLCs; with PSEN1-knockout affecting APP and TREM2 cleavage, and PSEN2-knockout altering endo-lysosomal biology.

Our observation of divergent effects of PSEN1 versus PSEN2-knockout support data from rodent models [44], where *Psen1* knockout mice are embryonic lethal [18,19] and *Psen2* knockout mice have a much milder phenotype [20]. Lethality of Psen1-knockout has been suggested to be down to impaired Notch signalling, and together with the finding of unchanged App processing in *Psen2* knockout mice [20], suggest that Psen1-associated γ-secretase is critical for these plasma-membrane cleavage events [45]. In contrast, PSEN2 has been shown to be enriched at the endo-lysosomal compartment [11,14,15], and the finding here of altered levels endo-lysosomal protein expression supports a major role for PSEN2 in this system. Indeed, endo-lysosomal effects of PSEN2 knockout and PSEN2-associated fAD mutations have also been described recently in mouse models [46].

The major role of PSEN1 in APP processing is supported in patients, as those with *PSEN1* mutations have an earlier age at onset compared to people carrying a *PSEN2* mutation [47,48]. Although both PSEN1 and PSEN2 fAD-linked mutations converge on altered APP processing, the later age at onset of PSEN2-associated mutations is consistent with PSEN1-containing γ-secretase contributing to 65% of overall Aβ production [23,37], consistent with a smaller overall impact on APP processing in PSEN2 knockout neurons. Frameshift mutations in *PSEN2* have been associated with fAD [49]; the same is not true for *PSEN1, NCSTN* and *PSENEN*, where loss-of-function is associated with familial hidradenitis suppurativa.

In a previous study which also aimed to compare PSEN1 and PSEN2 function in iPSC-neurons, Watanabe et al used a conditional knockout system to overcome developmental effects of double knockouts [15]. In contrast to the data presented here, they found that only double presenilin knockout lines had altered Aβ40 production and lines without PSEN2 had reduced Aβ42 production. These data were mirrored by an accumulation of APP C-terminal fragments only when both PSEN1 and PSEN2 were knocked out. Similar findings were presented in mouse N2A cell lines, where only the absence of both Psen1 and Psen2 affected Aβ generation and APP C-terminal fragment accumulation [50]. Further work is required to understand these disparate findings in these studies; however, it is conceivable that cells have a different compensatory capacity for constitutive and conditional loss of gene function – impacting redundancy for homologous proteins. Indeed, this has important implications for developmental versus ageing effects as well as the acute consequences of γ-secretase inhibitors.

It is intriguing that PSEN2 is upregulated in PSEN1-knockout neurons, suggestive of a compensatory response. This is relevant to fAD pathomechanisms, as PSEN2-associated γ-secretase has been shown to specify longer, disease-associated Aβ peptides [10,15]. It is thus conceivable that PSEN2 compensation in PSEN1-associated fAD can contribute to raised Aβ42:40 ratios. Indeed, we recently observed increased *PSEN2* expression in PSEN1-associated fAD iPSC-associated astrocytes [26], supporting a compensation in mutation-carrying cells as well as PSEN1-knockout models. We also described reduced Notch signalling in *PSEN1* mutation carrying neurons, mirroring PSEN1 loss-of-function [51]. However, it is crucial to appreciate that mutations in *PSEN1*/*PSEN2* are distinct from these loss-of-function experiments. There are clearly differing cellular phenotypes in *PSEN1* mutation cells and *PSEN1* knockout cells [23], as well as an established clinical distinction (fAD versus familial hidradenitis suppurativa).

A detailed understanding of the diversity of γ-secretase in a context-specific manner is critical for ongoing therapeutic discovery. γ-secretase inhibitor trials were halted due to adverse effects [16] (assumed to relate to effects on other substrates). Targeting of specific subclasses of γ-secretase, therefore, offers an attractive strategy to reduce unwanted off-target effects, a strategy for γ-secretase modulators/stabilisers as well as inhibitors [17].

## Limitations

Loss-of-function studies are informative for physiological functions of proteins, however, results can be hard to interpret when functional redundancy exists. The data presented supports divergent functions of the two catalytic subunits of γ-secretase. It is critical to distinguish these findings from fAD-associated mutations, which do not act through simple loss-of-function [23]. Indeed, this is evident from Aβ measurements in this study (Fig 2). Another limitation relates to Aβ quantification, where PSEN2-knockout has little effect on Aβ generation. Previous work suggests that PSEN2 is responsible for intracellular Aβ generation [14], and future work should quantify the effect of PSEN1/2 loss-of-function on the intracellular pool of Aβ.

## Conclusion

We describe divergent consequences of PSEN1 and PSEN2-knockout in iPSC-derived neurons and microglia. The data support a major role for PSEN1 in cleavage of membrane proteins such as APP and TREM2, whereas loss of PSEN2 function affects the endo-lysosomal system – mirroring the subcellular localisation of the two proteins. These findings suggest a therapeutic rationale for γ-secretase subtype-specific targeting, a strategy that could reduce wider effects of γ-secretase inhibition.

## Acknowledgements

We acknowledge the contribution of UCL Genomics for their support with karyotype analysis via whole genome sequencing (RRID number SCR_027010).

## Funding

This project was supported by a Pump Priming Award from ARUK London Network, enabling the generation of PSEN2-knockout iPSCs. CA was supported by a fellowship from Alzheimer’s Society (AS-JF-18-008). SW was supported by a senior fellowship from ARUK (ARUKSRF-2016B). LB and CA were supported by Race Against Dementia. FT was supported by EMBO Grant no. 9297. This work was supported by the National Institute for Health and Care Research University College London Hospitals Biomedical Research Centre.

## Author contribution

Investigation: CA, MBF, LB, FT,

Conceptualisation: CA, CLV, SW

Writing – review and editing: all authors

Funding acquisition: CA, MBF, SW

## Notes

### Competing Interest Statement

The authors have declared no competing interest.

