## Supplementary material for "Divergent consequences of PSEN1 knockout and PSEN2 knockout in stem cell derived models of the brain": Suppl Fig 1

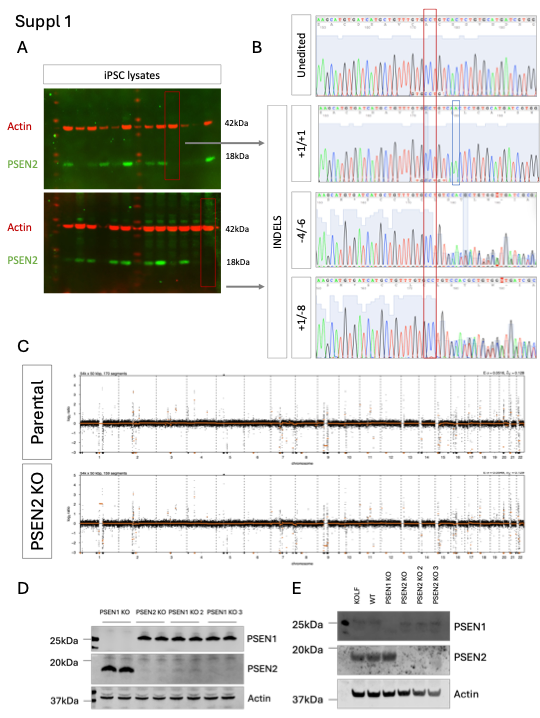

**Supplementary Figure 1. Quality control of newly generated PSEN2-knockout cells.**

A) Selection of leads clones via a Western blotting screen in individual iPSC colonies post genome editing. B) Sanger sequencing to confirm frameshift mutations in three lead clones, compared to the unedited parental line, and deconvolution of traces to detect indels via TIDE software [24] (red rectangle represents the PAM sequence and blue rectangle represents inserted base pair). C) Low coverage whole genome sequencing supports stable a karyotype in the lead PSEN2-knockout clone. Western blotting in D) iPSCs and E) neurons demonstrates absence of PSEN2 protein in all 3 PSEN2 knockout clones.
